# ABC-transporter activity and autocrine eicosanoid-signaling are required for germ cell migration a basal chordate

**DOI:** 10.1101/469098

**Authors:** Susannah H. Kassmer, Delany Rodriguez, Anthony DeTomaso

## Abstract

In the colonial ascidian *Botryllus schlosseri*, long-lived germline stem cells (GSCs) migrate to new germline niches as they develop during repetitive cycles of asexual reproduction. ABC-transporters are involved in the export of lipid-signaling molecules, but their roles in germ cell migration are poorly understood. Here, we show that in *Botryllus, abcc1* and *abcb1* are highly expressed in germ cells, and inhibition of ABC-transporter activity leads to failure of germ cell migration. Phospholipase A2 (PLA2) produces arachidonic acid, which is further metabolized to eicosanoid signaling molecules. In humans, 12-lipoxygenase (LOX) metabolizes arachidonic acid to12-Hydroxyeicosatetraenoic acid (12-S-HETE), which stimulates migration of mammalian cancer cells and smooth muscle cells. We show that PLA2 and LOX activity are required for germ cell migration. A potential homolog to the human receptor for 12-S-HETE, *BSgpr31*, is expressed in germ cells. 12-S-HETE rescues migration towards S1P in the presence of inhibitors of ABCC1, ABCB1, PLA2 or LOX, and a gradient of 12-S-HETE enhances chemotaxis towards S1P and stimulates motility. We conclude that 12-S-HETE is a secondary chemoattractant exported by ACB-transporters that is required for migration of germ cells towards S1P. We also find that in the presence of S1P, detection of an 12-S-HETE gradient initiates an autologous positive feedback loop that may sustain migration. This is the first report of an eicosanoid-signaling molecule regulating germ cell migration.

## Introduction

Cell migration is a fundamental process of development and maintenance of multicellular organisms and mediates tissue organization, organogenesis, immune response and homeostasis ^1^. Regulation of cell migration requires a complex interplay of signaling cascades that influence cell adhesion, polarization and cell motility. Temporal-spatial cues are tightly controlled, as dysregulation of cell migration can have catastrophic consequences for the organism, including developmental defects and cancer.

In many species, germ cells that are specified during embryonic development need to migrate across the embryo to reach the somatic gonad, where they will develop into eggs and testes ^2^. Germ cell migration is studied in a variety of organisms, and many features are widely conserved. Most germ cells undergo an active migration toward their somatic niche that is often be guided by a combination of attractive and repulsive cues ^3^.

In addition to being cell membrane components, research in the last two decades has shown that many lipids, now termed “bioactive lipids”, have critical cell signaling functions ^4^. Some lipid classes such as lysophospholipids (including sphingosine-1-phosphate), eicosanoids (e.g. prostaglandins), and endocannaboids can signal through receptors in the cell membrane ^4^. These lipids activate G-protein coupled receptors (GPCRs) of the Gi, Gq, and G12/13 type can be activated by gradients of bioactive lipids and thus influence cell polarity. Bioactive lipids associated with cell polarity include lysophospholipids (LPLs) and phosphatidylinositolphosphates (PIPs). LPLs are bioactive lipids that can be generated by hydrolytic cleavage of fatty acid from glycerophospholipids, a reaction catalyzed by phospholipases. Distinct phospholipases cleave off either one of the two (PLA1 and PLA2) fatty acid residues. The biosynthesis of eicosanoids is initiated by the activation of PLA2, leading to the release of arachidonic acid. Arachidonic acid is metabolized to a variety of eicosanoid signaling molecules such as thromboxanes, and leukotrienes ^4^.

So far, very little is known about the roles of bioactive lipids in germ cell migration. A potential role for lipid signaling in germ cells was first discovered in Drosophila, in which mutations in the lipid phosphate phosphatases (LPP), wunen and wunen2, disrupt directed migration of germ cells to the gonad ^3,5^, but the ligand has not been identified. However, this mechanism appears to be conserved, as LPPs also repel germ cells away from nearby somites in the zebrafish ^6^. Recently, our group discovered that migration of germ cells is directed by the lysophospholipid Sphingosine-1-phosphate (S1P) in the colonial ascidian *Botryllus schlosseri* ^7^.

*Botryllus* is a unique model to study germline biology, because an individual does not grow by increasing in size, but rather by a lifelong, recurring asexual budding process (called blastogenesis) during which entire bodies including all somatic and germline tissues. are regenerated de novo (Figure 1 A, B). This results in a constantly expanding colony of genetically identical individuals, called zooids, which are linked by a common extracorporeal vasculature (Figure 1).

**Figure 1:**
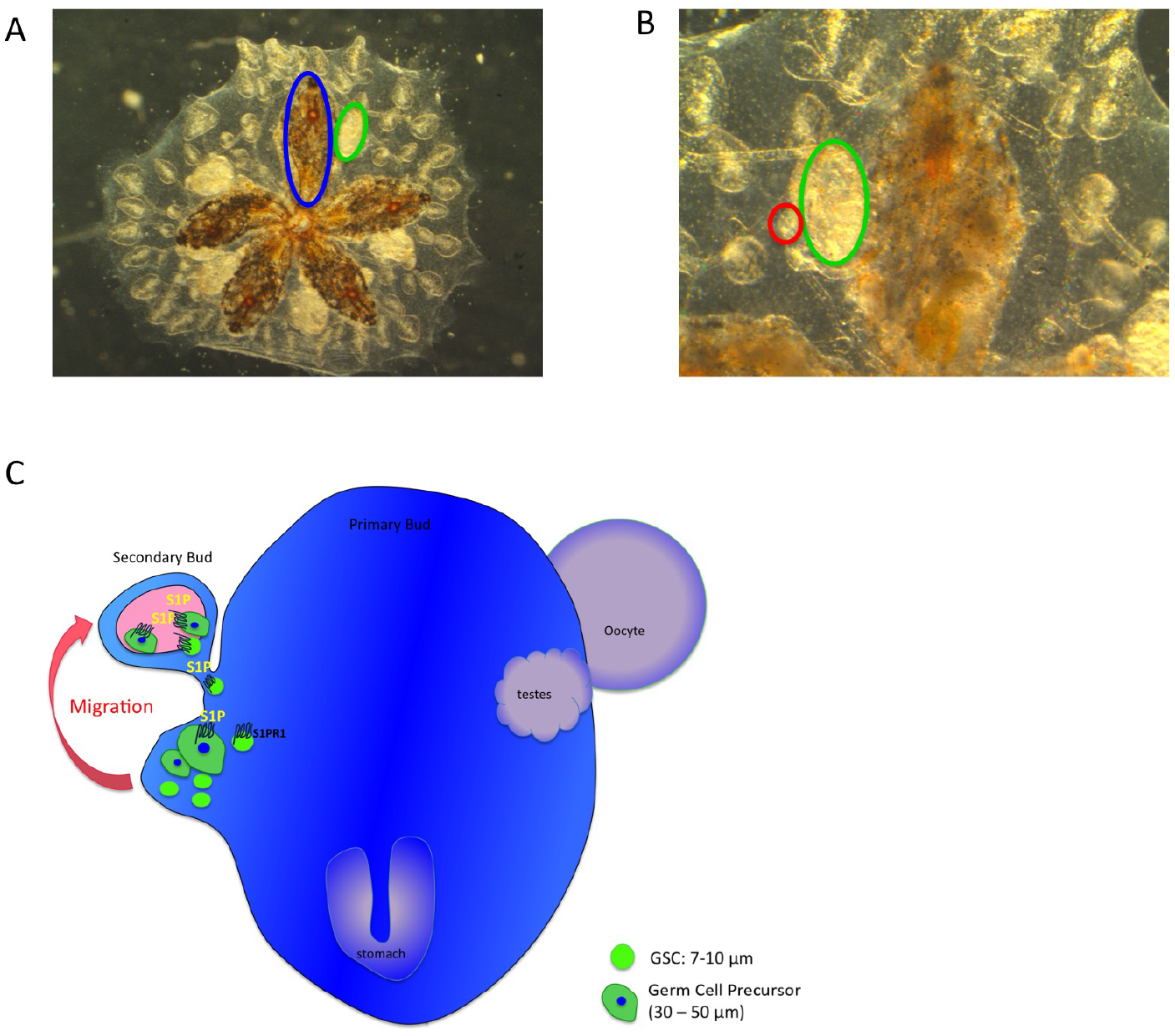
Morphology of Botryllus schlosseri colony, germ cell migration to secondary buds. A and B: colony morohology and asexual reproduction. All individual bodies within the colony are embedded in an extracellular matrix known as the tunic and share an extracorporeal vasculature. Adult zooids (blue outline) asexually reproduce by giving rise to primary (green outline) and secondary buds (red outline). C: Migration of germ cells to secondary buds. Secondary buds begin as small protrusions of the body wall of primary buds, and later form a closed double vesicle. This vesicle later grows and undergoes invaginations and tissue differentiation, completing development into the adult form. Germline stem cells and (GSC, 7-10μM) and germ cell precursors (30-50μM) migrate from the primary bud into the secondary bud at the time when the double vesicle forms. When the double vesicle is fully formed, germ cells have completed migration. Migration into the secondary bud is directed by a chemotactic gradient of sphingosine-1-phosphate, which is secreted within the secondary bud and detected by sphingosine-1-phosphate-receptor-1 expressed by the migrating germ cells (Kassmer et al 2015).

The best understood regenerative process in *Botryllus* occurs in the germline. Like most metazoans, *Botryllus* sets aside a population of PGCs early in embryogenesis that are responsible for gametogenesis for the life of the organism ^8^. However, unlike most model organisms, *Botryllus* retains a population of mobile, self-renewing, lineage-restricted adult germline stem cells (GSCs) that retain pluripotency for life of the individual ^9^. During the weekly development of new zooids, a new germline niche is formed, and GSCs migrate from the old germline niche to the new. A subset of these GSCs settle, differentiate into gametes (zooids are hermaphrodites and make sperm and eggs), while others self-renew and migrate to the next generation. Asexual development is synchronized throughout the colony, and migration occurs over a defined 48h period as GSCs leave the primary bud niche and enter the secondary bud niche via the vasculature joining the two (Figure 1A, B, C). During this window, GSCs are also found in the colony vasculature and ampullae. GSC migration is controlled by a S1P gradient, which is synthesized in the new niche and binds to the GPCR S1PR1on migrating GSCs ^7^(Figure 1C).

The ATP binding cassette (ABC) are transport proteins that are conserved in all phyla from prokaryotes to humans ^10^. ABC transporters shuttle a variety of hydrophobic lipophilic compounds across the cell membrane in an ATP-dependent manner, including bioactive lipids such as S1P, leukotrienes and prostaglandins ^11^. Some ABC transporters play roles in cell migration. In *Drosophila*, a germ cell attractant is geranylgeranylated and secreted by mesodermal cells in a signal peptide-independent manner through an ABCB transporter of the MDR family ^12^. Chemotaxis of human dendritic cells to CCL19 requires stimulation with the exogenous leukotriene C4, an eicosanoid transported out of the cell via ABCC1 ^13^.

Cells responding to a primary chemoattractant can secrete secondary chemoattractants that increase the robustness of the primary chemotactic response ^14,15^. Neutrophils release leukotriene B_4_ (LTB_4_) to enhance their chemotactic response to the inflammatory cue fMLP. Binding of fMLP to cell surface receptors initiates leukotriene biosynthesis by stimulating the conversion of arachidonic acid (AA) to LTB_4_. LTB_4_ is released as a secondary chemoattractant, and stimulates neutrophil motility through its interaction with its cognate receptor BLT1. The LTB_4_ is packaged in exosomes, which are secreted in a polarized fashion to the region of the cell with the highest fMLP concentration, setting up a gradient along the cell itself. Failure to form or detect the secondary chemoattractant causes an impaired chemotactic response ^15^.

In the present study, we show that activity of ABCC1 and ABCB1 is required for directed migration of *Botryllus* germ cells in shallow gradients of the primary chemoattractant S1P *in vitro* and for homing towards developing germline niches *in vivo*. We identify the lipoxygenase-product 12-S-HETE as a secondary chemoattractant that enhances chemotaxis towards shallow gradients of S1P. *Botryllus* germ cells express a homolog of the human 12-S-HETE receptor GPR31. We conclude that shallow gradients of S1P induce activation of LOX and production and ABC-transporter-mediated export of 12-S-HETE, resulting in autocrine stimulation of chemotaxis.

## Results

### *Vasa*-positive cells express ABCC1 and ABCB1

We have shown previously that *vasa*-positive germline stem cells can be prospectively isolated from the blood of *Botryllus schlosseri* by flow cytometry, using a mononclonal antibody against Integrin-alpha-6 (IA6) ^7^. By quantitative real time PCR, *abcc1* and *abcb1* mRNA are highly enriched in IA6+ cells, with *abcb1* is expressed at higher levels than *abcc1* (6.2 fold enrichment vs 2.2 fold enrichment, Figure 2A). We confirmed expression of *abcc1* and *abcb1* in *vasa*-positive cells migrating to secondary buds (Arrows, Figure 2B) by fluorescent in situ hybridization (FISH).

**Figure 2:**
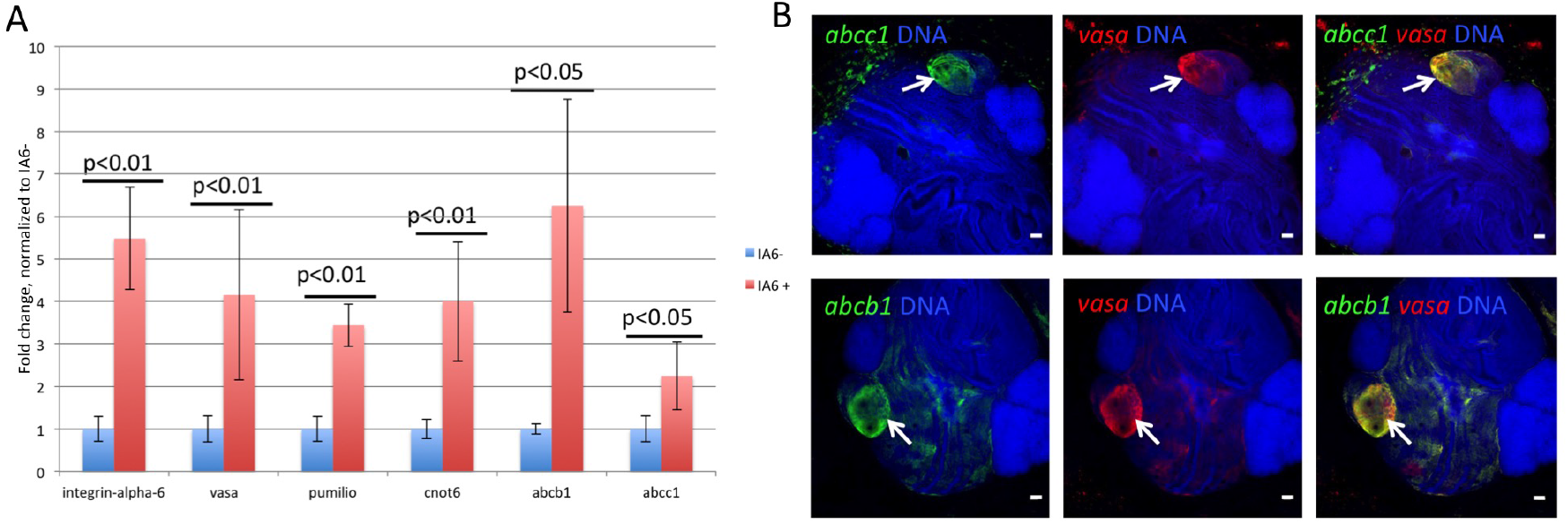
Botryllus germ cells express *abcb1* and *abcc1*. A: *Abcb1* and *abcc1* are expressed in Integrin-alpha-6-positive (IA6+) germline stem cells. IA6-positive and- negative cells were isolated by flow cytometry and expression of germ cell marker genes (*vasa, pumilio, cnot6*) and ABC-transporters (*abcb1* and *abcc1*) was assessed by quantitative real time PCR. Relative quantification was performed using the 2^-ΔΔCT^-method, with *actin* as control gene. Data are expressed as averages of the relative expression ratio (fold change), normalized to IA6-negative cells. Standard deviations were calculated for each average expression ratio (n=3). Statistical analysis was performed using Student’s *t*-test. B: Representative examples of double-labeled fluorescent in situ hybridization showing expression of *abcc1* and *abcb1* in *vasa*-positive cells. All *vasa*-positive (red) germ cell precursors (arrows) co-express *abcc1 and abcb1* (green, arrows). Red and green channels are shown individually with nuclear counterstaining (blue), and merged images on the right show co-expression of both genes (yellow) Scale bars = 20μm.

### ABCB1 and ABCC1 activity is required for migration towards low concentrations of S1P

To test whether inhibition of ABC transporter activity affects migration of *Botryllus* germ cells we isolated IA6+ cells by flow cytometry and assessed their migratory activity to S1P in our transwell migration assay.

In the presence of inhibitors of either ABCC1 or ABCB1, migratory activity to a low concentration of S1P (0.2μM) is significantly reduced (p≤ 0.05, figure 3A). An inhibitor of both ABCC1 and ABCB1 reduces migration even further, almost to control levels. In contrast, migration in the presence of a high concentration of S1P (2μM) is not significantly affected by ABC-transporter inhibition (Figure 3A).

**Figure 3:**
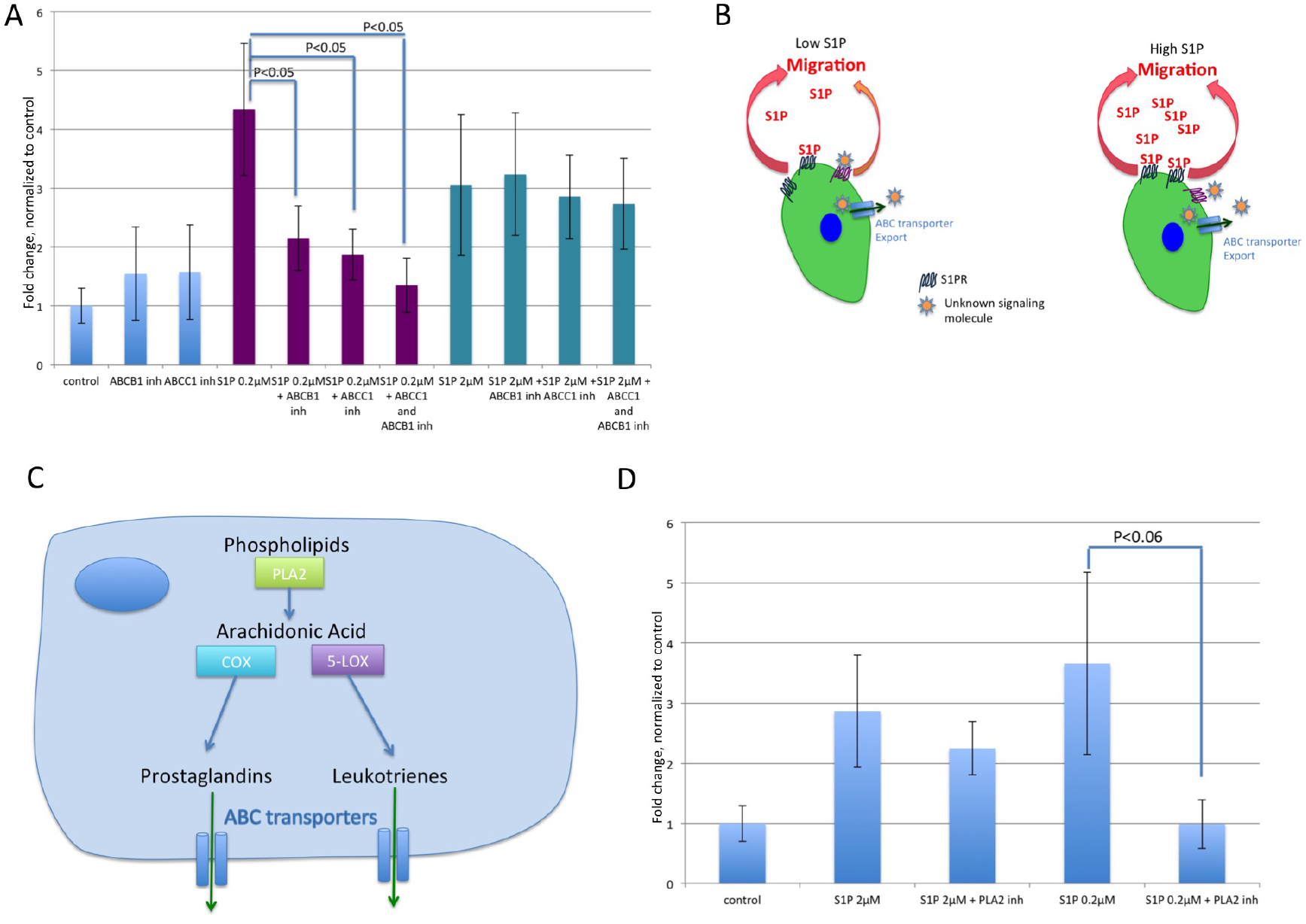
Migration of Integrin-alpha-6-positive germ cell precursors towards low concentrations of sphingosine-1-phosphate (S1P) depends on ABC-transporter activity. A) Migration assay of IA6-positive cells in response to different concentrations of S1P, with or without inhibitors of ABC-transporters, as indicated. No S1P was added to control wells. IA6+ cells were added to the upper chamber of a transwell system coated with laminin, and after 2h, migrated cells in the lower chamber were counted. Data are expressed as fold changes of numbers of migrated cells, normalized to unstimulated controls (n=4). Statistical analysis was performed using Student’s *t*-test. B: Hypothetical model of ABC-transporter mediated export of an unknown secondary chemoattractant signaling molecule. Migration to shallow gradients of S1P depends on ABC-transporter activity. Low-level stimulation of the S1P-receptor might induce production and ABC-transporter mediated secretion of an unknown signaling molecule, which provides a secondary chemoattractant signal to enhance migration towards S1P. Steeper gradients of S1P are sufficient to stimulate chemotaxis, and ABC-transporter-mediated export of a secondary chemoattractant is not required. C: The cytoplasmic enzyme phospholipase A2 (PLA2) generates arachidonic acid from phospholipids. Arachidonic acid is further metabolized by either lipoxygenases (LOX) or Cyclooxygenases (COX) to generate bioactive lipids such as leukotrienes or prostaglandins, which are exported out of the cytoplasm by ABC transporters. D: migration assay of IA6+ cells in response to S1P in the presence of an inhibitor of PLA2. Inhibition of PLA2 completely blocks the migratory activity to 0.2uM of S1P, but has no significant effect on migration towards 2uM of S1P. Data are expressed as fold changes of numbers of migrated cells, normalized to unstimulated controls (n=4). Statistical analysis was performed using Student’s *t*-test.

These results suggest that ABCC1 and ABCB1 might export a secondary chemoattractant that enhances chemotaxis in the presence of low concentrations of the primary chemoattractant S1P (Figure 3B). In the presence of high concentrations of S1P, stimulation through the S1P receptor alone is sufficient to stimulate chemotaxis.

### Phospholipase A2 activity and lipoxygenase activity are required for migration to S1P

Next, we aimed to assess for a possible secondary chemoattractant that might be exported by ABCC1 and ABCB1. ABC transporters export a variety of substrates. Among these are a variety of lipid signaling molecules, such as phospholipids and derivatives of arachidonic acid ^11,16^. In humans, the cytoplasmic enzyme phospholipase A2 (PLA2, Fig3C) generates the polyunsaturated omega-6 fatty acid arachidonic acid from phospholipids. Arachidonic acid is further metabolized by either lipoxygenases (LOX) or Cyclooxygenases (COX) to generate bioactive lipids such as leukotrienes or prostaglandins, which are exported out of the cytoplasm by ABC transporters (Fig3C) ^16^. To test whether a derivative of arachidonic acid plays a role in germ cell migration towards low concentrations of S1P, we assessed migratory activity in the presence of an inhibitor of PLA2. Inhibition of PLA2 completely blocked the migratory response to 0.2μM of S1P (Fig3D), but only had a mild effect on migration to a high concentration of S1P (2μM, Fig3D). These results show that a derivative of arachidonic acid is required for the migratory response to low concentrations of S1P. To assess whether this derivative is a product of cyclooxygenase or lipoxygenase, we tested the migratory response to 0.2μM of S1P in the presence of several inhibitors of COX-1 or COX-2 or an inhibitor of lipoxygenases. *Botryllus schlosseri* lipoxygenase (*bslox*) has homology to 5-LOX, 12-LOX and 15-LOX from different organisms, and *bslox* expression is enriched in IA6+ cells (Figure 6A, fig S1C)). An inhibitor that blocks the activity of all 3 human lipoxygenases blocks germ cell migration to 0.2μM of S1P, whereas COX-inhibitors had no effect (Fig 4A). Furthermore, cyclooxygenases are not expressed at significant levels in IA6+ germ cells (Fig S1C). Migration to a high concentration of S1P (2μM) is not affected by LOX inhibition (Fig4A). These results show that an arachidonic-acid-derived product of lipoxygenase is exported by ABCC1 or ABCB1 and is required for migration towards low concentrations of S1P.

**Figure 4:**
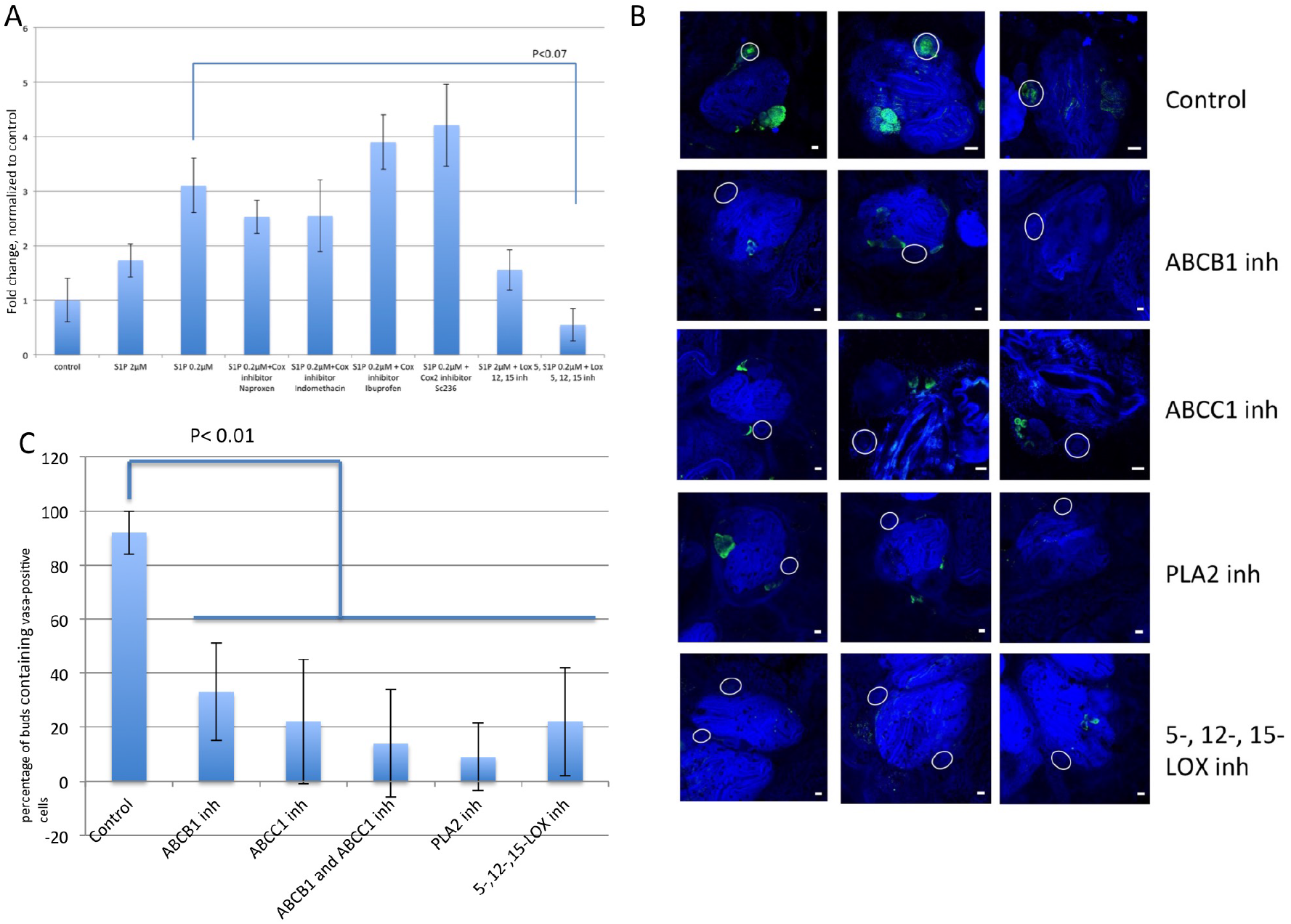
Migration towards low concentrations of S1P depends on activity of Lipoxygenase. A: Migration assay of IA6+ cells in response to S1P, with or without inhibitors Cox1, Cox2 or 5-,12-, and 15-LOX, as indicated. Data are expressed as fold changes of numbers of migrated cells, normalized to unstimulated controls (n=4). Statistical analysis was performed using Student’s *t*-test. B and C: Homing of germ cells in vivo depends on activity of PLA2 and LOX. Animals were treated with inhibitors of ABC transporters, PLA2 or 5-,12-, and 15-LOX for 3 days, starting at stage A1, and fixed at stage B2, when the secondary bud forms a closed double vesicle (n=4). Controls were left untreated. *Vasa*-FISH was performed on fixed animals, and the number of secondary buds containing germ cells were counted by confocal microscopy. Nuclei were counterstained with Hoechst 33342 (blue). Scale bars = 20μm. In control animals, *vasa*-positive germ cells (green) homed into the double vesicle stage secondary buds (circles). All 4 inhibitors significantly reduced migration of vasa-positive cells to secondary buds. Graph in C shows the percentage of double vesicle stage secondary buds containing *vasa*-positive cells for each treatment. Error bars represent the standard deviation for each average (n=4). Statistical analysis was performed using Student’s *t*-test.

### Inhibition of ABCC1, ABCB1, PLA2 or Lipoxygenases reduces migration of germ cells to secondary bud niches *in vivo*

To test whether migration of germ cells to secondary buds in vivo requires activity of ABCC1, ABCB1, PLA2 or Lipoxygenase, we allowed *B. schlosseri* colonies to develop in the presence of inhibitors. Drugs were added to the seawater at stage A1, when the new generation of zooids first begin to develop. Following the migration period, when the GSCs have migrated from old to new germline niches (Figure 1C), the colonies where fixed in formaldehyde and germ cell migration analyzed by vasa-FISH (Figure 4B). *Vasa*-positive cells in the new niche were quantified using confocal microscopy. In colonies treated with inhibitors, significantly fewer vasa-positive cells (green) entered into new germline niche (white circles) (p≤0,01, Figure 4C) compared to vehicle treated controls.

### Botryllus germ cells express a receptor for the 12-LOX product 12-S-HETE

Different types of lipoxygenases use arachidonic acid as a substrate to generate a variety of downstream signaling molecules, including leukotrienes, lipoxins, and other fatty acids such as different types of hydroperoxyeiocatetraenoic acid (HPETE) and hydroxyicosatetraenoic acid (HETE) (Figure 5A). To assess which lipoxygenase-product is responsible for enhancing migration to S1P, we tested an inhibitor of 5-LOX, as well as a cysteinyl leukotriene receptor antagonist. Neither of these significantly affected migration to S1P (Figure 5B). These data suggested that a product of either 12-LOX or 15-LOX is responsible for migration to S1P. 12-LOX generates 12-S-HETE, a signaling molecule that stimulates migration of cancer cells and smooth muscle cells ^17^. In humans, the G-protein-coupled receptor GPR31 is a high affinity receptor for 12-S-HETE ^18^. We identified a *Botryllus* homolog of GPR31 in our EST database, and have named it *Bsgpr31*. In humans, another receptor for 12-S-HETE, and for the 15-LOX product 15-S-HETE is Leukotriene B4 receptor 2 ^19^, but we were not able to identify a homolog of this receptor in *Botryllus*. Using qPCR, we found that *Bsgpr31* expression is significantly enriched in IA6+ cells and expressed at levels comparable to *vasa* (FigS1C and D). We also identified a *Botryllus* homolog of the closely related GPCR “Trapped in endoderm” *tre-1*, a fatty acid receptor that is important for germ cell guidance in *Drosophila* ^20^. This receptor is not expressed in Integrin-alpha-6-positive cells (Fig S1C and D). Using in situ hybridization, we confirmed that *Bsgpr31* mRNA is expressed exclusively in *vasa*-positive cells migrating to secondary buds (Figure 5C, white arrows). Expression of *abcc1, abcb1* or *gpr31* does not show any significant changes during the blastogenic cycle in fertile and infertile animals in our published transcriptomes (Fig S2)^21^, indicating that these genes are always expressed in germ cells. This is in line with our own observation that migratory activity of IA6+ cells in vitro is independent of the blastogenic stage of the original colony.

**Figure 5:**
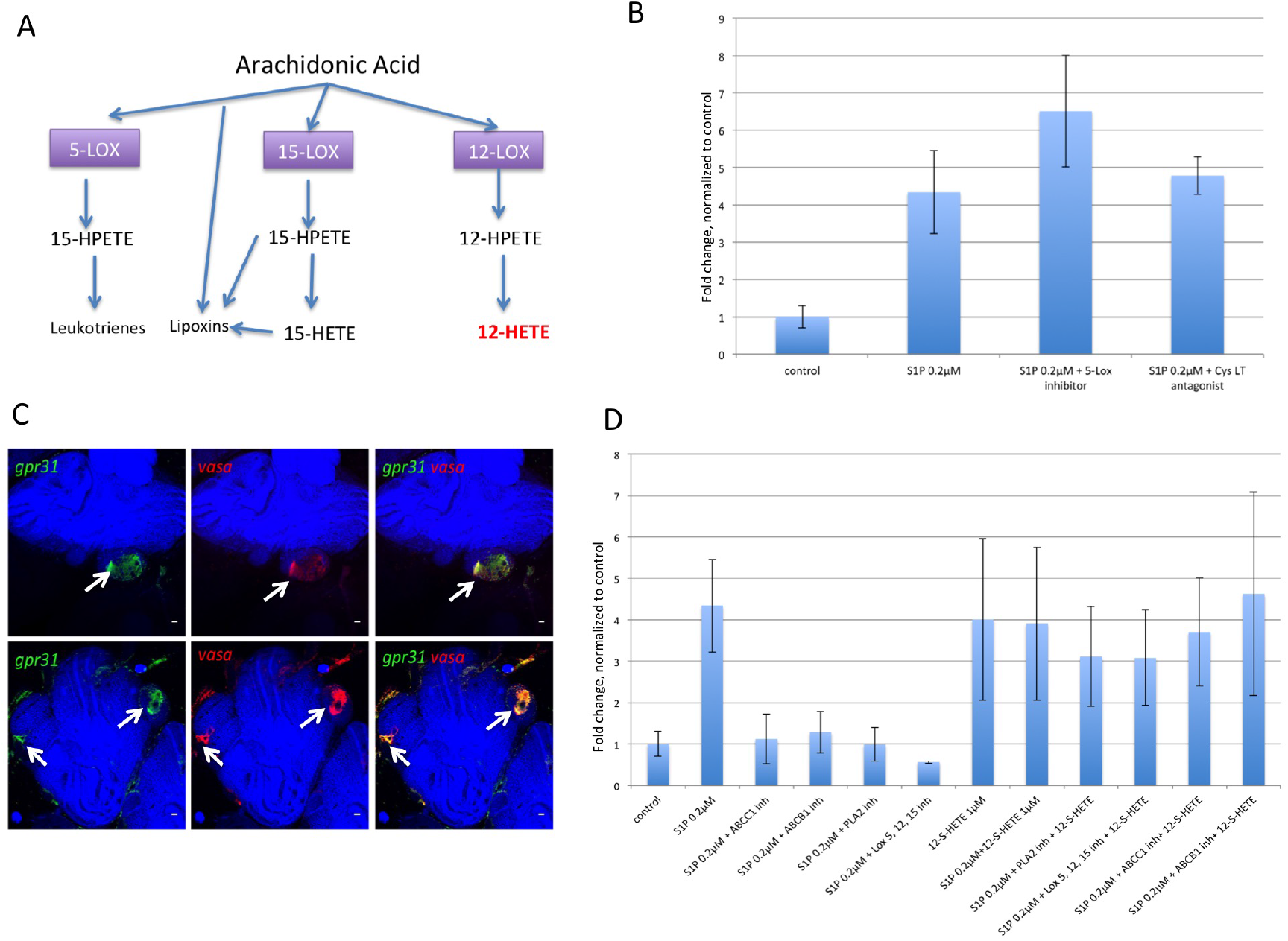
Botryllus germ cells express the 12-S-HETE receptor gpr31, 12-S-HETE rescues migration in the presence of ABC-transporter and lipoxygenase inhibitors. A: Migration assay of IA6-positive cells in response to 0.2μM S1P, with or without inhibitors 5-LOX or Cysteinyl Leukotriene Receptor, as indicated. Data are expressed as fold changes of numbers of migrated cells, normalized to unstimulated controls (n=4). Statistical analysis was performed using Student’s *t*-test. B: In humans, 3 different types of lipoxygenase metabolize arachidonic acid to various signaling agents. 12-Lox metabolizes arachidonic acid to 12(S)-hydroperoxy-5Z,8Z,10E,14Z-eicosatetraenoic acid (12(S)-HpETE). 12(S)-HpETE is rapidly reduced to 12(S)-HETE by ubiquitous cellular peroxidases, such as Glutathione peroxidases. C: Representative examples of double-labeled fluorescent in situ hybridization showing expression of *gpr31* in *vasa*-positive cells. All *vasa*-positive (red) germ cell precursors (arrows) co-express *gpr31* (green, arrows). Red and green channels are shown individually with nuclear counterstaining (blue), and merged images on the right show co-expression of both genes (yellow) Scale bars = 20μm. D: Migration assay of IA6-positive cells in response to 0.2μM S1P and/or 12-S-HETE, with or without inhibitors of ABC-transporters, PLA2 or 5-,12-, and 15-LOX, as indicated. Data are expressed as fold changes of numbers of migrated cells, normalized to unstimulated controls (n=4). Statistical analysis was performed using Student’s *t*-test.

### The 12-LOX product 12-S-HETE stimulates germ cell migration and rescues inhibition of ABC-transporters, PLA2 and lipoxygenases

As *vasa*-positive cells in *Botryllus* express the putative receptor for 12-S-HETE, Bsgpr31, we aimed to test whether 12-S-HETE would stimulate migratory activity of IA6+ cells *in vitro*. 12-S-HETE alone has a stimulating effect on migration of IA6+ cells (Figure 5D). 12-S-HETE induces migration nearly to nearly the same extent as 0.2μM S1P. In combination, both molecules in combination induce the same amount of migration as 12-S-HETE alone. Importantly, adding exogenous 12-S-HETE rescues migratory activity to 0.2μM S1P in the presence of inhibitors of ABCC1, ABCB1, PLA2 or LOX (Figure 5D). These data suggest that migration to 0.2μM S1P is dependent on the secondary chemoattractant-12-S-HETE, and also demonstrates that the inhibitors used are not affecting chemotaxis non-specifically.

### 12-S-HETE acts as a secondary chemoattractant and increases chemotaxis to S1P

To test the effect of 12-S-HETE on S1P-induced chemotaxis, we characterized the migratory behavior of cells exposed to a chemotactic gradient in a 3D matrix. The chemotactic gradient was established by adding S1P and/or 12-S-HETE to the left reservoir of a chemotaxis chamber, and filtered seawater to the right reservoir, and cells were unmanipulated, or exposed to inhibitors of endogenous 12-S-HETE synthesis. We initially used identical concentrations of S1P and 12-S-HETE to those used in the transwell assays, and both the direction and average distance traveled by the GSCs under each condition are shown in Figure 6.

**Figure 6:**
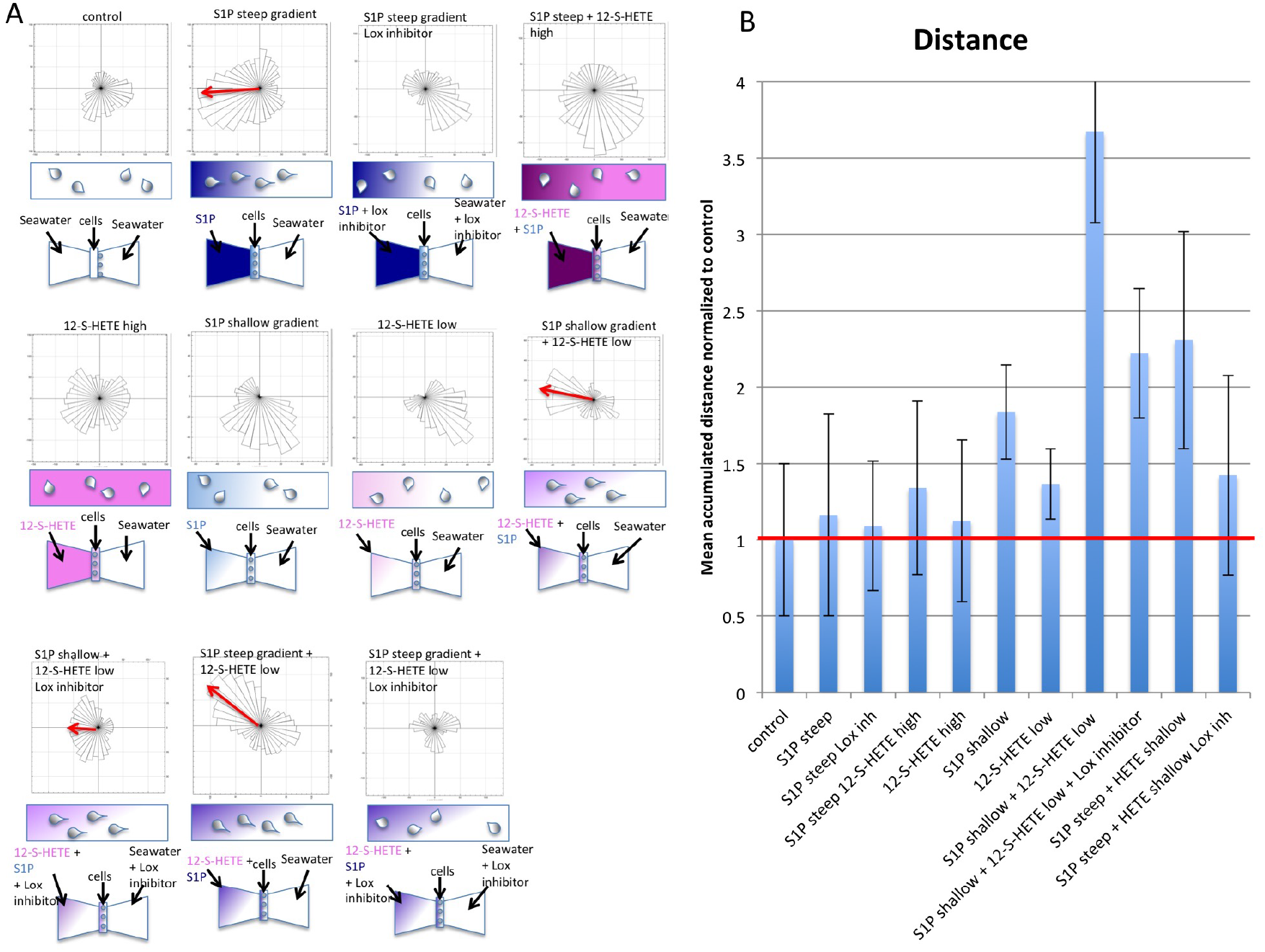
Migration to low concentrations of S1P depends on 12-S-HETE production: A: Chemotaxis assay. Chemotaxis was analyzed by live imaging of cells embedded in Matrigel in a chemotaxis chamber. The right reservoir contained filtered seawater, and for steep gradients the left reservoir contained 500nM 12-S-HETE (pink) or 0.2μM S1P (blue) or both. For shallow gradients, 50nM 12-S-HETE (light pink) or 0.2μM S1P (light blue) (final concentrations) were added to the indicated corner of the left reservoir. For controls, both reservoirs contained filtered seawater. Data from 3 independent experiments were combined and plotted as rose diagrams showing the directionality of cell paths for each condition tested. B: Average accumulated distance for cells migrating in each condition (n=3), normalized to unstimulated controls, with standard deviation. The red line indicates the distance migrated by control cells (unstimulated).

Initially, we characterized the response when a high concentration of S1P was added to the left reservoir (steep gradient, Figure 6A). Under these conditions, cells directionally migrate towards the left side (Figure 6A), and cover more distance then unstimulated controls (Figure 6B). This response required endogenous 12-S-HETE production, as inhibition of Lipoxygenase abolishes S1P-directed chemotaxis, and also reduced the total distance travelled (Figure 6B).

When 12-S-HETE and S1P are both added to the reservoir, cells cover more distance than in S1P alone, but loose directionality (Figure 6A, B). When 12-S-HETE alone was added to the left reservoir, cells migrate randomly (Figure 6A) and cover more distance than unstimulated controls (Figure 6B). We hypothesize that these results are due to the fact that arachidonic acid derived eicosanoids are very small molecules that diffuse rapidly, and upon release, the resulting gradients are shallow and transient. Cells that secrete such secondary chemoattractants must use a mechanism to create short-range gradients alone the cell axis ^22^. In neutrophils responding to a primary gradient of fMLP, the secondary gradient is formed by localized secretion of LTB_4_ in exosomes to the region of the cell experiencing the highest concentration of fMLP. Our data suggest that when 12-S-HETE is added to the left reservoir it diffuses too quickly for a stable gradient to form, and the cells are exposed to 12-S-HETE from all sides and lose their ability to perform directional migration towards the primary chemoattractant S1P. Thus it appears that 12-S-HETE stimulates migration non-directionally if not in a gradient along the cell boundary.

We next attempted to create a shallow gradient of 12-S-HETE in the chemotaxis chamber by adding a very low concentration of 12-S-HETE to one of the corners of the left reservoir (pre-filled with seawater) immediately before live-imaging, so that 12-S-HETE would diffuse more slowly towards the cells. Under these conditions (12-S-HETE shallow gradient), 12-S-HETE alone does not induce chemotaxis, but cells cover more distance than controls (Figure 6A, B). We next used the same technique to create a very shallow gradient of S1P. In this very shallow S1P gradient, cells cover more distance compared to controls, and even compared to cells migrating in a steep S1P gradient, but they are not able to migrate directionally towards S1P (Figure 6A, B). It has been shown that in suboptimal concentrations of chemoattractant, cells turn more frequently ^23^, potentially facilitating the search for regions with optimal concentrations of chemoattractant. These results suggest that very low concentrations of S1P are not sufficient to stimulate production and secretion of 12-S-HETE.

However, when we used the technique described above to add a combined shallow gradient of 12-S-HETE and S1P to the corner of the reservoir, the cells were able to migrate directionally towards S1P, and covered more distance than in any of the other conditions tested (Figure 6A, B).

This demonstrates that when present as a gradient, 12-S-HETE enhances chemotaxis to the primary chemoattractant S1P and acts as a secondary chemoattractant in *Botryllus* germ cells (illustrated in Figure 7A). Together, this data suggests that under normal conditions, detection of S1P induces 12-S-HETE secretion in a polarized manner, and that this signal relay mechanism is required for chemotaxis under physiological S1P concentrations. We next wondered if chemotaxis in a shallow exogenously applied gradient of S1P and 12-S-HETE also required autocrine production of 12-S-HETE. When a lipoxygenase inhibitor is added, chemotaxis and distance are reduced, indicating that autocrine production of 12-S-HETE is required for chemotaxis in cells migrating towards S1P (illustrated in Figure 7B), even in the presence of an external gradient of 12-S-HETE. This suggests the presence of a positive feedback loop, where autocrine production of 12-S-HETE is induced by extracellular gradients of S1P and 12-S-HETE (illustrated in Figure 7A).

**Figure 7:**
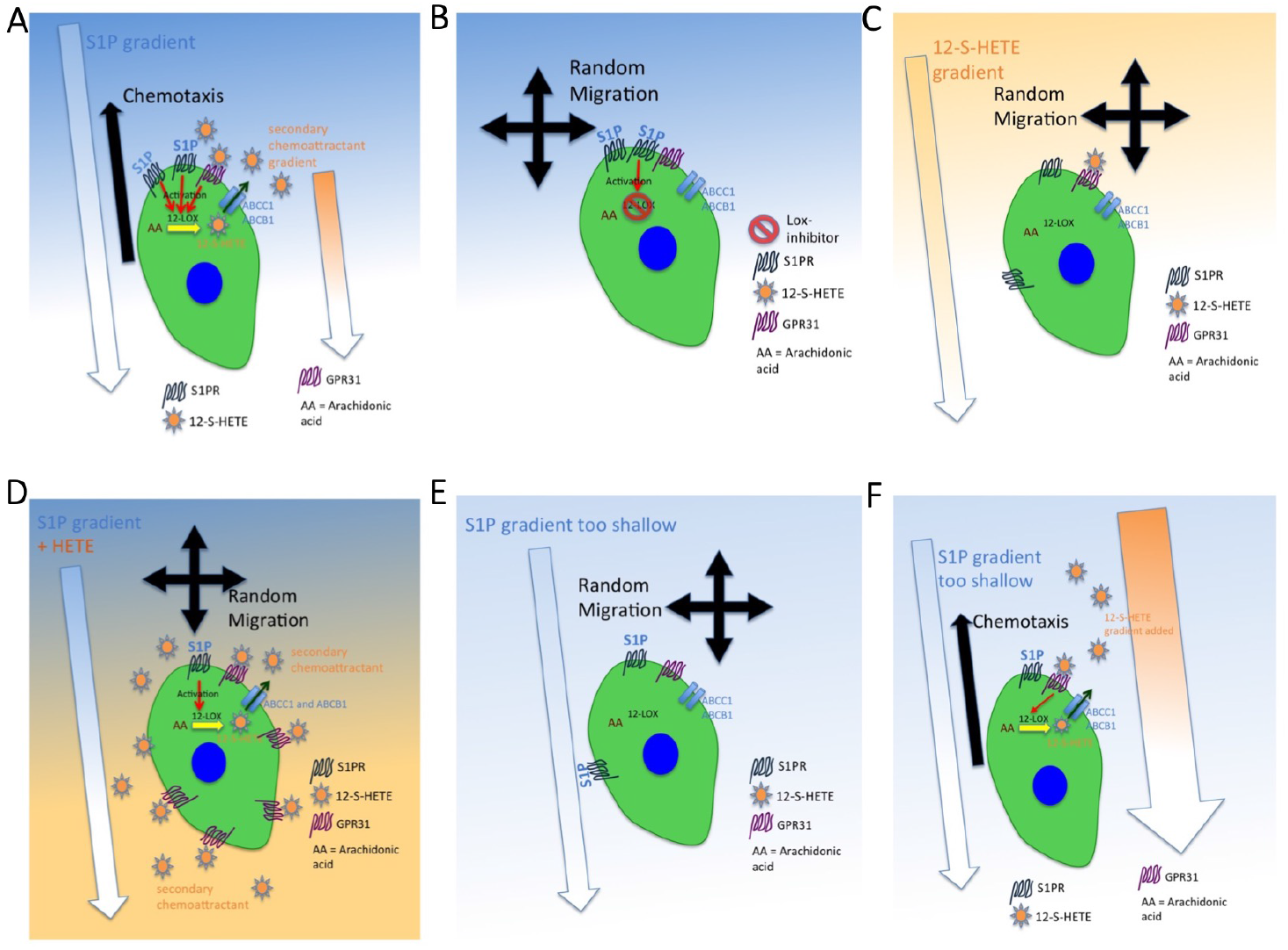
Hypothesized mechanism for 12-S-HETE acting a secondary chemoattractant during migration of germ cells towards S1P. A: In a gradient of S1P, S1P receptors are activated and stimulate activity of 12-LOX, leading to production and export of 12-S-HETE. 12-S-HETE acts as a secondary chemoattractant, and binds to GPR31 on the cell surface, resulting in enhanced chemotaxis. Activation of the 12-S-HETE-receptor GPR31 stimulates activity of 12-LOX, sustaining chemotaxis. B: When 12-LOX is inhibited, no 12-S-HETE is secreted, resulting in lack of chemotaxis towards S1P and random migration. C: A gradient of 12-S-HETE alone does not stimulate chemotaxis, because the primary chemoattractant is missing. D: When 12-S-HETE is present in uniform concentration in all directions, cells lose their directionality, resulting in random migration. E: In sub-optimal concentrations of S1P or a gradient that is too shallow, S1P receptors are not sufficiently activated to stimulate secretion of 12-S-HETE. F: When an artificial gradient of 12-S-HETE is added to a shallow gradient of S1P, chemotaxis towards S1P is stimulated. Activation of the 12-S-HETE-receptor GPR31 stimulates activity of 12-LOX, sustaining chemotaxis.

Together, this data suggests that under normal conditions, detection of S1P induces 12-S-HETE secretion in a localized manner, and that this signal relay mechanism is required for chemotaxis under physiological S1P concentrations. Given that results using a high concentration of 12-S-HETE suggested that this molecule diffuses rapidly, we next wondered if chemotaxis in a shallow exogenously applied gradient of 12-S-HETE also required downstream, autocrine 12-S-HETE stimulation. In the next set of experiments cells treated with Lipoxygenase inhibitor were assayed under the same conditions (shallow S1P and shallow 12-S-HETE), and under these conditions chemotaxis and distance are reduced, indicating that autocrine production of 12-S-HETE is required for chemotaxis in cells migrating towards S1P (Figure 6; Illustrated in Figure 7B), even in the presence of an external shallow gradient of 12-S-HETE. This suggests the presence of a positive feedback loop, where autocrine production and localized secretion of 12-S-HETE is induced by extracellular gradients of S1P and 12-S-HETE (illustrated in Figure 7A).

Interestingly, the same results were observed when a shallow gradient of 12-S-HETE is combined with a high concentration of S1P: chemotaxis towards S1P is enhanced and mean migration distance increases when combined with a shallow 12-S-HETE gradient. (Figure 6; illustrated in Figure 7A). However even under these high S1P concentrations, inhibition of lipoxygenase reduces chemotaxis and distance in this experiment, providing further evidence that an autocrine-feedback loop regulates 12-S-HETE secretion and may function to sustain chemotaxis in cells migrating towards S1P (illustrated in Figure 7A).

## Discussion

This is the first report of an eicosanoid signaling molecule directing germ cell migration, and the first report of 12-S-HETE-stimulated migration of germ cells. We show that 12-S-HETE acts as a secondary chemoattractant that facilitates chemotaxis of germ cells towards the primary chemoattractant S1P. This mechanism is termed signal relay, and it can extend the spatial range over which cells can be directed in a primary gradient. In exponential gradients, signal relay could potentially attract cells in areas where the slope of the primary gradient is shallow. For example, LTB_4_ enables directed migration of neutrophils when the primary gradient is too shallow to induce chemotaxis ^15^. We show that migration of *Botryllus* germ cells towards low concentrations of S1P requires the activity of ABC-transporters and lipoxygenase (Illustration in Figure 7A, results in Figure 4, 5D and 6). Inhibition of LOX abolishes the ability of cells to directionally migrate towards S1P (Illustration in Figure 7A, results in Figure 6A). In a very shallow gradient of S1P, cells can only perform directed migration when a gradient of the 12-LOX product 12-S-HETE is present (Illustration in Figure 7E and F, results in Figure 6A), indicating that 12-S-HETE is a secondary chemoattractant that is required for directional migration towards the primary chemoattractant S1P. This idea is further supported by the fact that a gradient of 12-S-HETE alone does not induce chemotaxis (Illustration in Figure 7C, results in Figure 6).

Chemotaxis towards S1P is blocked by inhibition of Lox, even in the presence of an externally added gradient of 12-S-HETE, suggesting the presence of a positive feedback loop, where autocrine production of 12-S-HETE is induced by extracellular gradients of S1P and 12-S-HETE (illustrated in Figure 7A and F). When a higher concentration of 12-S-HETE is added directly to one of the reservoirs of the chemotaxis chamber, cells migrate randomly (Illustration in Figure 7D), suggesting that a small molecule such as 12-S-HETE diffuses too quickly to form a stable gradient along the cell axis. The same phenomenon has been reported for other small fatty acid secondary chemoattractants, such as leukotriene B4 (LTB_4_) ^22^. Therefore, we hypothesize that the export and release of 12-S-HETE by migrating germ cells must be spatially and temporally controlled. In human neutrophils migrating towards a primary chemoattractant, secreted LTB_4_ is packaged in exosomes ^22^ By gradually releasing LTB_4_, exosomes may prevent LTB_4_ profiles from rapidly reaching saturating concentrations ^15^. If 12-S-HETE is packaged in such exosomes, ABC transporters might be involved in controlling secretion of 12-S-HETE from exosomes. Alternatively, 12-S-HETE secretion could be spatially controlled along the axis of the migrating cell, and occur only at the leading edge, to form a gradient along the cell axis.

There is one main difference between chemotaxis in the 3D matrix and the transwell migration assay: there can be no stable gradient of 12-S-HETE in the transwell – this small molecule diffuses too quickly, and, therefore, the cells are likely sensing 12-S-HETE from all directions. This may explain why there was no additive effect of S1P and 12-S-HETE in the transwell migration assay (Figure 5D), but in the 3D matrix, cells perform more chemotaxis and cover more distance in the presence of a gradient of 12-S-HETE when it is added to a gradient of S1P (Figure 6 A, B). In general, migration in a 3D matrix is more physiological than migration on a 2D plastic surface, so the results from the IBID assay are likely mimicking the in vivo situation more closely.

While ABC transporters play known roles in dendritic cell and cancer cell migration ^16^, only one study has so far reported a requirement of an ABC transporter in germ cell migration. MDR49 regulates the export of farnesyl-modified mating factors in yeast and is expressed in the drosophila mesoderm. MDR49 mutants have defects in PGC migration ^12^. Here, we show that in *Botryllus*, germ cell migration depends on activity of ABCB1 and ABCC1 and the export of an eicosanoid secondary chemoattractant. This is the first study suggesting a role for signal relay and secretion of a secondary chemoattractant in germ cells migrating in a primary chemotactic gradient. In addition, our data suggests that once a 12-S-HETE gradient is detected, cells amplify this signal via continued secretion to the region of the cell experiencing the highest concentration, initiating a positive feedback loop that would help drive and sustain chemotaxis. Polarized secretion could be due to redistribution of enzymes responsible for 12-S-HETE production, or the ABC transporters required for secretion.

The roles of bioactive lipids in germ cell migration are poorly understood, but there is growing evidence for their importance. In Drosophila germ cell migration, the GPCR Tre1 directs migration through the midgut ^20^. While the ligand for Tre1 still unknown, the closest mammalian homolog, GPR84, binds to medium chain fatty acids. In *Drosophila* and zebrafish, lipid phosphate phosphatases are required for directed migration of germ cells ^5,6^. To our knowledge, no other studies have addressed the role of lipids in germ cell migration in other species. Here we show that the bioactive lipids S1P and 12-S-HETE regulate germ cell migration in an invertebrate chordate, suggesting that bioactive lipids may play conserved roles in directing germ cell migration across phyla.

12-S-HETE has been shown to stimulate cell migration in human cell types. Specifically, it induces migration of cancer cells on laminin ^24^. This is relevant since *Botryllus* germ cells also migrate on laminin ^7^. 12-S-HETE induces PKC-dependent cytoskeletal rearrangements in tumor cells, resulting in increased motility ^17^ and stimulates aortic smooth muscle cell migration ^25^. Interestingly, upregulation of 12-LOX induces a migratory phenotype in cancer cells ^26^, suggesting autocrine stimulation by 12-S-HETE plays a role in metastasis. In a carcinoma cell line, activation of beta-4 integrin induces translocation of 12-LOX to the membrane and upregulates its enzymatic activity ^27^. Finally, both 12-S-HETE and GPR31 expression positively correlate to prostrate cancer grade and progression in humans (Honn et al., 2016). In context of these observations, we hypothesize that in *Botryllus* germ cells, S1P-signaling and binding of Integrin-alpha-6 to laminin might induce translocation of 12-LOX or ABC trannsporters to the leading edge, resulting in localized secretion of 12-S-HETE.

When a small amount of 12-S-HETE is added to a very shallow gradient of S1P, cells cover considerably more distance than with S1P alone (Figure 6B), suggesting that at the right concentration, 12-S-HETE not only enhances directional migration but also increases motility. Cells undergoing directional migration require environmental guidance cues as well as the ability to initiate and sustain motility. Depending on the organism, migrating germ cells must sustain directed migration for twenty-four to forty-eight hours ^3^. In *Botryllus*, migration of germ cells to new, germline niches occurs during a defined 48h period ^28^. It is common for a migrating cell to require more than one signal to induce this type of directed migration and motility. In mice, the chemokine SDF-1 provides the guidance cue to migrating primordial germ cells, whereas signaling of SCF through the receptor c-kit enhances motility ^3^. Our results suggest that a 12-S-HETE positive feedback response may be responsible for maintaining directional migration.

In conclusion, migration of GSC towards a shallow gradient of S1P depends on ABC-transporter mediated export of a secondary chemoattractant produced by lipoxygenase. The 12-lipoxygenase-product 12-S-HETE acts as a secondary chemoattractant and enhances directional migration towards shallow gradients of the primary chemoattractant S1P. In turn, our data suggest that 12-S-HETE must also be in a gradient, and moreover that exogenous12-S-HETE stimulation induces endogenous production, amplifying the secondary signal. We have discovered a novel mechanism of signal relay required for germ cell chemotaxis. Signal relay had been previously studied in neutrophils and *Dictyostelium discoideum* ^29^, but to our knowledge has never been described in germ cells. Given the role of 12-S-HETE in cancer cell migration ^26^, our study suggests that a conserved eicosanoid based signal relay mechanisms might operate in many other cell types across different species.

## Materials and Methods

### Animals

*Botryllus schlosseri* colonies used in this study were lab-cultivated strains, spawned from animals collected in Santa Barbara, CA, and cultured in laboratory conditions at 18-20 °C according to ^30^. Colonies were developmentally staged according to ^31^.

### Cell Sorting

Genetically identical, stage matched animals were pooled, and a single cell suspension was generated by mechanical dissociation. Whole animals were minced and passed through 70 μm and 40 μm cell strainers in ice-cold sorting buffer (filtered sea-water with 2% horse serum and 50mM EDTA). Anti-Human/Mouse-CD49f–eFluor450 (Ebioscience, cloneGoH3) was added at a dilution of 1/50 and incubated on ice for 30 min and washed with sorting buffer. Fluorescence activated cell sorting (FACS) was performed using a FACSAria (BD Biosciences) cell sorter. Samples were gated IA6 (CD49f)-positive or – negative based on isotype control staining (RatIgG2A-isotype-control eFluor450, Ebioscience). Analysis was performed using FACSDiva software (BD Biosciences). Cells were sorted using a 70 μm nozzle and collected into sorting buffer.

### Quantitative RT PCR

Sorted cells were pelleted at 700g for 10min, and RNA was extracted using the Nucleospin RNA XS kit (Macherey Nagel), which included a DNAse treatment step. RNA was reverse transcribed into cDNA using random primers (Life Technologies) and Superscript II Reverse Transcriptase (Life Technologies). Quantitative RT-PCR (Q-PCR) was performed using a LightCycler 480 II (Roche) and LightCycler DNA Master SYBR Green I detection (Roche) according to the manufacturers instructions. The thermocycling profile was 5 min at 95, followed by 45 cycles of 95 °C for 10 sec, 60 °C for 10 sec. The specificity of each primer pair was determined by BLAST analysis (to human, *Ciona* and *Botryllus* genomes), by melting curve analysis and gel electrophoresis of the PCR product. To control for amplification of genomic DNA, ‘no RT’-controls were used. Primer pairs were analyzed for amplification efficiency using calibration dilution curves. All genes included in the analysis had CT values of <35. Primer sequences are listed in Supplemental table 1. Relative gene expression analysis was performed using the 2^-ΔΔCT^ Method. The CT of the target gene was normalized to the CT of the reference gene *actin* : ΔC_T_ = C_T (target)_ – C_T (actin)_. To calculate the normalized expression ratio, the ΔC_T_ of the test sample (IA6-positive cells) was first normalized to the ΔC_T_ of the calibrator sample (IA6-negative cells): ΔΔC_T_= ΔC_T(IA6-positive)_-ΔC_T(IA6-negative)_. Second, the expression ratio was calculated: 2^-ΔΔCT^= Normalized expression ratio. The result obtained is the fold increase (or decrease) of the target gene in the test samples relative to IA6-negative cells. Each qPCR was performed at least three times on cells from independent sorting experiments gene was analyzed in duplicate in each run. The ΔC_T_ between the target gene and *actin* was first calculated for each replicate and then averaged across replicates. The average ΔC_T_ for each target gene was then used to calculate the ΔΔC_T_ as described above. Data are expressed as averages of the normalized expression ratio (fold change). Standard deviations were calculated for each average normalized expression ratio (n=6). Statistical analysis was performed using Student’s T-test.

### *In Situ* Hybridization

Whole mount *in situ* hybridization was performed as described in ^32^. Briefly, *B. schlosseri* homologs of genes of interest were identified by tblastn searches of the *B. schlosseri* EST database (http://octopus.obs-vlfr.fr/public/botryllus/blast_botryllus.php) using human or Ciona (when available) protein sequences. Primer pairs were designed to amplify a 500-800 bp fragment of each transcript (Primer sequences in Supplemental table 1). PCR was performed with Advantage cDNA Polymerase (Clontech, 639105) and products were cloned into the pGEM-T Easy vector (Promega, A1360). *In vitro* transcription was performed with SP6 or T7 RNA polymerase (Roche, 10810274001, 10881767001) using either digoxigenin or dinitrophenol labeling. HRP-conjugated anti-digoxigenin antibody (Roche, 11207733910) or HRP-conjugated anti-dinitrophenol antibody (Perkin Elmer, FP1129) were used to detect labeled probes by fluorophore deposition (Fluorescein or Cyanine 3) using the TSA Plus System (Perkin Elmer, NEL753001KT). Nuclei were stained with Hoechst 33342 (Thermofisher). Imaging of labeled samples was performed using an Olympus FLV1000S Spectral Laser Scanning Confocal.

### Calcein assay of ABC transporter activity

Cells were isolated as described above for cell sorting and filtered through a 10uM cell strainer. Cells were incubated with Calcein-AM (1/1000, Thermo Fisher) in seawater for 30 minutes at room temperature in the presence of 10uM CP 1000356 hydrochloride (ABCB1-inhibitor), 10uM Probenecid (ABCB1 inhibitor) or 10uM Reversan (ABCB1 and ABCC1 inhibitor). Controls were incubated in Calcein only. Unstained controls were incubated in seawater only. Live cells were gated using forward and side scatter properties. Mean fluorescence intensity (MFI) of Calcein fluorescence was analyzed using a FACSAria (BD Biosciences) cell sorter and FACSDiva software (BD Biosciences).

### Transwell Migration Assay

Transwell filters with 8μm pore size inserted in a 24 well plate (Corning) were coated with laminin over night at 4°C and briefly air dried before adding 50,000 sorted cells, resuspended in 100μl filtered seawater with 10% DMEM and 1% FBS. The bottom of the well contained filtered seawater with 10% DMEM (Corning) /1% FBS (Corning) and 1% Primocin (InvivoGen). Sphingosine-1-phosphate (0.2 - 2μM, Echelon), 12(S)-Hydroxy- (5Z,8Z,10E,14Z)-eicosatetraenoic acid (80nM, Sigma-Aldrich) and 10uM CP 1000356 hydrochloride (ABCB1-inhibitor,), 10uM Probenecid (ABCB1 inhibitor), 10uM Reversan (ABCB1 and ABCC1 inhibitor), 10uM AACOCF3 (inhibitor of phospholipase A2), 0.5uM 2-TEDC (inhibitor of 5-,12- and 15-lipoxygenase), 10uM Zileuton (inhibitor of 5-lipoxygenase), 1uM BAY-u 9773 (Cysteinyl leukotriene receptor antagonist), 10uM (S)-(+)-Ibuprofen (Cox-1 inhibitor), 1mM SC 236 (Cox-2 inhibitor) (all from Tocris) were added to the bottom chamber as indicated. For controls, the bottom chamber contained filtered seawater with 10% DMEM /1% FBS and 1% Primocin. After 2 hours incubation at room temperature, nuclei in the bottom well were stained with Hoechst 33342 (1/1000, Thermofisher) and manually counted. All assays were performed in triplicates with cells from 4 independent sorts. Statistical analysis was performed using Student’s t-test.

### Small Molecule Inhibitor Treatment

*Botryllus* colonies were incubated in 5 ml of seawater containing 25μM Reversan (ABCB1 and ABCC1 inhibitor), 100μM Probenecid (ABCB1 inhibitor) 20μM CP 1000356 hydrochloride (ABCB1-inhibitor) or 25 μM 2-TEDC (inhibitor of 5-,12- and 15-lipoxygenase) or 14 μM AACOCF3 (inhibitor of phospholipase A2). Controls were incubated in seawater. Treatment was started at stage A2, and animals were fixed at stage B2 and analyzed by *in situ* hybridization as described above. Each treatment was performed on 3 genetically identical colonies. Buds containing *vasa*-positive cells were counted on each treated and untreated colony under an epifluoresence microscope. Statistical analysis was performed using Student’s t-test.

### Chemotaxis Assay

Cells were isolated from the blood of Botryllus colonies, filtered through 10μm cell strainers and embedded in 50% Matrigel (Corning) with 50% filtered seawater. Whole blood was used to avoid the stress of cell sorting, since only vasa+ cells expressing S1PR1 ^33^ or GPR31 (Figure 5C and S1C and D) will respond to stimulation with S1P or 12-S-HETE. 6μl of gel were added to each chamber of an ibidi μ-slide Chemotaxis (ibidi GmbH, Martinsried, Germany). The gel was allowed to polymerize in a humidified chamber for 30 minutes at room temperature before adding filtered seawater to the right reservoir. The left reservoir was filled with filtered seawater containing Sphingosine-1-phosphate (0.2μM, Echelon) or 12(S)-Hydroxy-(5Z,8Z,10E,14Z)-eicosatetraenoic acid (500nM, Sigma-Aldrich) or both. For samples containing 0.5uM 2-TEDC (inhibitor of 5-, 12- and 15-lipoxygenase), the inhibitor was added to both reservoirs. For controls, both reservoirs contained filtered seawater. To achieve a shallow gradient of S1P, 6μl of 2 μM S1P were added to the corner of the left reservoir containing 60μl of filtered seawater. To achieve a shallow gradient of 12-S-HETE, 6μl of 50nM 12-S-HETE were added to the corner of the left reservoir containing 60μl of filtered seawater. Live imaging was performed on a Leica SP8 confocal microscope at 15s intervals. Cell paths were tracked manually using the Manual Tracking Plugin in Image J. At least 30 cells were tracked in each field of view, and the data from 3 independent experiments were combined for the final analysis. Cell paths were analyzed using the Chemotaxis and Migration Tool Version 1.01 (https://ibidi.com/chemotaxis-analysis/171-chemotaxis-and-migration-tool.html) for Image J.

## Acknowledgements

We would like to thank Amro Hamdoun for helpful discussions. Ben Lopez at the MCDB and NRI microscopy core facility is acknowledged for help with live imaging.

