## supplemental information for "ABC-transporter activity and autocrine eicosanoid-signaling are required for germ cell migration a basal chordate"

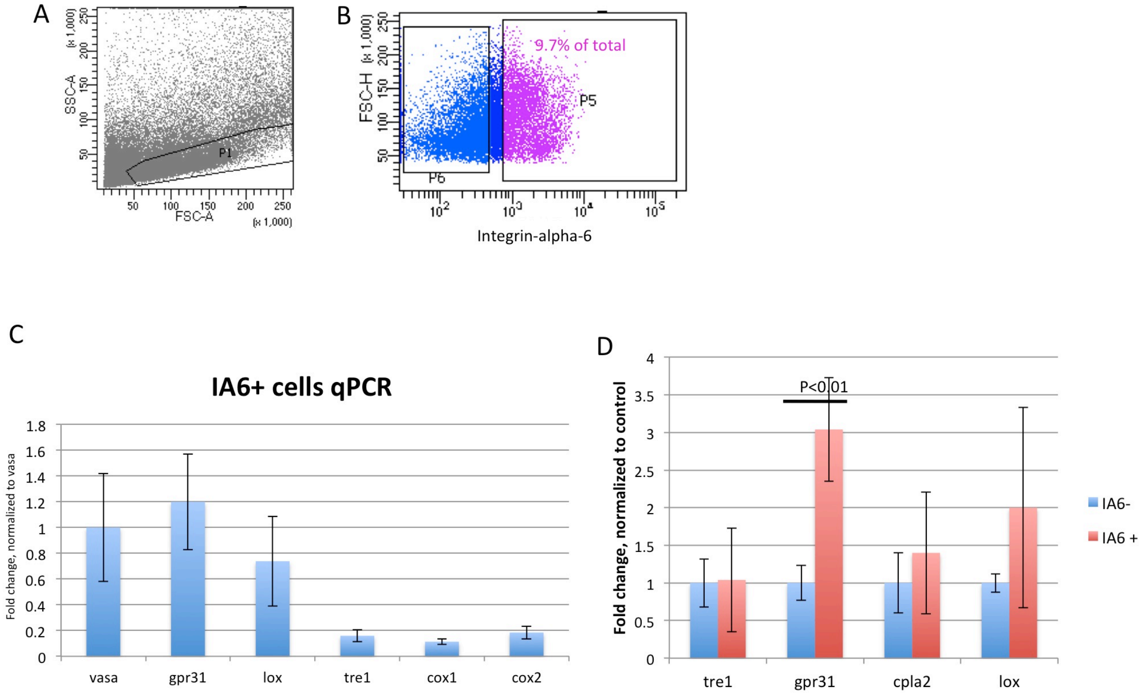


**Supplemental Figure 1**: A: Isolation of Integrin-alpha-6-(IA6) positive cells by flow cytometry. Forward-Side-Scatter gating on live cells. B: Cells within the live cell gate, plotted for forward scatter and Integrin-alpha-6-fluorescence. Gate around IA6- cells in p6 was determined using an isotype control antibody. IA6+ cells in p6 comprise about 9.7% of total cells. C: IA6+ cells were isolated by flow cytometry and expression of *vasa, gpr31, lox, tre1, cox1* and *cox2* were analyzed by quantitative real time PCR. Relative quantification was performed using the $2^{-\Delta\Delta CT}$-method. Data are expressed as averages of the relative expression ratio (fold change), normalized to *vasa*.. Standard deviations were calculated for each average expression ratio (n=4). D: IA6-positive and- negative cells were isolated by flow cytometry and expression of *tre1, gpr31, cytosolic phospholipase a2, cpla2*) and ABC-transporters (*abcb1* and *abcc1*) was assessed by quantitative real time PCR. Relative quantification was performed using the $2^{-\Delta\Delta CT}$-method, with *actin* as control gene. Data are expressed as averages of the relative expression ratio (fold change), normalized to IA6-negative cells. Standard deviations were calculated for each average expression ratio (n=4). Statistical analysis was performed using Student's *t*-test (n=4).

### GPR31

| | infertile | fertile |
|---|---|---|
| A1 | 1 | 1 |
| A2 | 0.780692072 | 0.858131578 |
| B1 | 0.874530347 | 0.945237859 |
| B2 | 0.831144248 | 0.637873041 |
| C1 | 0.958803171 | 0.978669569 |
| C2 | 0.940888823 | 1.314125211 |
| D | 1.197872223 | 0.848157835 |

### ABCB1

| | infertile | fertile |
|---|---|---|
| A1 | 1 | 1 |
| A2 | 1.333605418 | 0.978136839 |
| B1 | 1.431100243 | 1.15140182 |
| B2 | 1.168386987 | 1.006248996 |
| C1 | 1.429030308 | 1.042679405 |
| C2 | 1.586021971 | 1.19967918 |
| D | 1.176567763 | 1.252461052 |

### ABCC1

| | infertile | fertile |
|---|---|---|
| A1 | 1 | 1 |
| A2 | 1.115297028 | 0.980482778 |
| B1 | 2.080704778 | 1.177504138 |
| B2 | 1.756696106 | 1.122607086 |
| C1 | 1.447558821 | 1.045940944 |
| C2 | 1.351838892 | 0.74168011 |
| D | 0.972279195 | 0.972558007 |

**Supplemental Figure 2**: mRNA-seq analysis of changes in gene expression of grp31, abcb1 and abcc1 during the blastogenic cycle and in fertile vs infertile colonies. Tables on the left show fold changes of differential expression for each gene normalized to blastogenic stage A1. Tables on the right show fold changes of differential expression for each gene in fertile animals normalized to infertile animals. mRNA seq analysis was performed at each stage of the blastogenic cycle (A1, A2, B1, B2, C1, C2 and D) on a total of 3 fertile genotypes and 3 infertile genotypes (Rodriguez, Sanders et al. 2014). After Quality Control analysis, the sequences were mapped to our publicly available Botryllus schlosseri EST database Bot_asmb assembly (04.05.2011, A. Gracey) (consisting of 50,107 contigs and representing several genotypes, both fertile but non-pregnant and infertile, at different stages of the blastogenic cycle. In order to identify putative homologs of the ESTs in our database, we performed a translated BLAST (blastx) analysis using the non-redundant human protein database (NCBI version 4/25/13). Differential expression analysis was performed with DESeq 1.10.1 using triplicates for the analysis.

**Supplemental Methods:**

**bsgpr31 cloning:**
Using tBLASTn, we searched the Botryllus-EST database using the human GPR31 protein sequence and identified a 2340 bp sequence that shows some homology to GPR31: Score = 29.6 bits (65), Expect = 3.5, Identities = 24/115 (20%), Positives = 44/115 (38%), Gaps = 1/115 (0%). In NCBI-BLASTX, this sequence shows homology to GPR36 (platynereis dumerii), melatonin receptor (Ciona) 5-hydroxytryptamine receptor (Ciona) and GPR31 (platynereis dumerii). When aligned directly with human GPR31, there is significant overlap: Score 29,6, Query cover 38%, E-value 2e-04.
Primers:
Gpr31 cloning forward ACGGAACCCAGCATACAGTG
Gpr31 cloning reverse CATTCCCGGTATTCTCGCCA
Product size 608

**qPCR Primers: Product size 140-180bp**

Gpr31 qPCR forward TGAAAGATGACTCGTCGCCC
GPR31 qPCR reverse CTACAAGAGATCGGCGGCTT

Vasa qPCR forward GGCGGATTTAGCGATGATGAG
Vasa qPCR reverse TTCCCCCATAGCGACTGTTAGAC

Pumilio qPCR forward GTCCATGTACGGTTCTGCCA
Pumilio qPCR reverse TTCGGGAAACGGTTGTTCCT

Cnot6 qPCR forward CTACGACGACACTGAGCAGG
Cnot6 qPCR reverse TCGGTGGGGGCATCCTAATA

Integrin-alpha-6 qPCR forward ACTTCCGGCACGAACAAGAT
Integrin-alpha-6 qPCR reverse GTACAACAGGGTAACCGGGG

Abcc1 qPCR forward TGAGGTTCCCGATTTCCGTG
Abcc1 qPCR reverse CGTCTTCCACGACGATAGCA

Abcb1 qPCR forward CGCCAATTAGGACGCCAAAG
Abcb1 qPCR reverse GACTTACGGCCTGGCTTTCT

Lox qPCR forward TCATGTATTGCGCCGACTGT
Lox qPCR reverse AACATCTTGGCCAGCATCCA

Pla2 qPCR forward AGGTTATCACGGGGATCGGA
Pla2 qPCR reverse CGGATTTTCGGTCGGGAGAA

Tre1 qPCR forward GACCATGAACGCGATGCAAA
Tre1 qPCR reverse CGGCAAAAGCATCGTCGTAG

**Cloning primers for FISH probes:**

Abcc1 cloning forward TACGTTCTGGTGGTTCACGG
Abcc1 cloning reverse GCGTACAGCTTGAGGACCTT
Product size 668

Gpr31 cloning forward ACGGAACCCAGCATACAGTG
Gpr31 cloning reverse CATTCCCGGTATTCTCGCCA
Product size 608

Vasa cloning forward AGGCACTATGATTCAGCCTGTG
Vasa cloning reverse ATCATAATCACCCGTCTCGCG
Product size 976

Abcb1 cloning forward ACGCAGTGAGAAGATTGCCA
Abcb1 cloning reverse ACCGAGCAGAATTTCACGGA
Product size 742
